# Characterization of peptide-protein relationships in protein ambiguity groups via bipartite graphs

**DOI:** 10.1101/2021.07.28.454128

**Authors:** Karin Schork, Michael Turewicz, Julian Uszkoreit, Jörg Rahnenführer, Martin Eisenacher

**Affiliations:** Medizinisches Proteom-Center, Medical Faculty, Ruhr-University Bochum, Bochum, Germany; Medical Proteome Analysis, Center for Protein Diagnostics (PRODI), Ruhr-University Bochum, Bochum, Germany; Department of Statistics, TU Dortmund University, Dortmund, Germany

**Author notes:** Institute for Clinical Biochemistry and Pathobiochemistry, German Diabetes Center (DDZ), Leibniz Center for Diabetes Research at the Heinrich Heine University Düsseldorf, Düsseldorf, Germany. German Center for Diabetes Research (DZD), Partner Düsseldorf, München-Neuherberg, Germany. These authors contributed equally to this work (Jörg Rahnenführer, Martin Eisenacher). Corresponding author. (ME).

## Abstract

In bottom-up proteomics, proteins are enzymatically digested into peptides before measurement with mass spectrometry. The relationship between proteins and their corresponding peptides can be represented by bipartite graphs. We conduct a comprehensive analysis of bipartite graphs using quantified peptides from measured data sets as well as theoretical peptides from an *in silico* digestion of the corresponding complete taxonomic protein sequence databases. The aim of this study is to characterize and structure the different types of graphs that occur and to compare them between data sets. We observed a large influence of the accepted minimum peptide length during in *silico* digestion. When changing from theoretical peptides to measured ones, the graph structures are subject to two opposite effects. On the one hand, the graphs based on measured peptides are on average smaller and less complex compared to graphs using theoretical peptides. On the other hand, the proportion of protein nodes without unique peptides, which are a complicated case for protein inference and quantification, is considerably larger for measured data. Additionally, the proportion of graphs containing at least one protein node without unique peptides rises when going from database to quantitative level. The fraction of shared peptides and proteins without unique peptides as well as the complexity and size of the graphs highly depends on the data set and organism. Large differences between the structures of bipartite peptide-protein graphs have been observed between database and quantitative level as well as between analyzed species. In the analyzed measured data sets, the proportion of protein nodes without unique peptides ranged from 6.4% to 55.0%. This highlights the need for novel methods that can quantify proteins without unique peptides. The knowledge about the structure of the bipartite peptide-protein graphs gained in this study will be useful for the development of such algorithms.

## Introduction

The digestion of intact proteins to peptides via enzymes like trypsin is a requirement for high-throughput bottom-up proteomics based on mass spectrometry (MS) [1–3]. Because of this, peptides are identified and quantified directly from the MS measurements instead of proteins. The protein ambiguity problem describes the challenge to build a list of proteins for which evidence exists that they are present in the respective sample using protein inference methods [4, 5]. Many identified peptides cannot unambiguously be assigned to a single protein, as the respective peptide sequence is part of multiple protein sequences in the underlying database. Because of shared peptides there is the need to form protein ambiguity groups, which consist of proteins whose presence in a sample cannot be completely decided using the peptides identified in the sample. Beyond this, for protein quantification peptide quantities need to be summarized to protein quantities. This is often done by using only unique peptides, although including shared peptides would be desirable to exploit the full potential of the data and to avoid inaccuracies [6–8]. The worst case, a valid quantification of proteins without any quantified unique peptide, is very challenging and not solved yet despite the large variety of different protein quantification methods available [9, 10].

To illustrate and facilitate the protein inference and quantification steps, bipartite graph representations of the protein-peptide relationships have been used [11–15]. A bipartite graph is an undirected graph *G* (with node set *N*(*G*) and edge set *E*(*G*)) whose nodes are grouped into two sets *N*_1_(*G*) and *N*_2_(*G*), so that each edge connects a node of the first set with a node of the second set. In the context of protein-peptide-relationships, proteins are represented by nodes from *N*_1_(*G*) and peptides by nodes from *N*_2_(*G*). One straightforward approach would be to add an edge if and only if the considered peptide sequence is part of the protein sequence based on database knowledge. Another possibility is to draw an edge only if the corresponding peptide was quantified in a specific data set (see section *Construction of bipartite graphs*). As not all proteins are connected to each other via chains of shared peptides, the bipartite graph for the whole data set can be divided into smaller connected components. These components can be analyzed and handled separately during protein inference or quantification.

The structure of the connected components is determined by the underlying protein sequence database and contains valuable information on how difficult the protein inference or quantification problem is in this specific case. The M-shaped example in Fig 1A has two protein nodes that each have one unique peptide node and there is a shared peptide node connecting both protein nodes. In this case, the protein inference step is quite simple, as both proteins are likely to be present in the underlying samples because unique peptides exists. For protein quantification however, the shared peptides contain valuable information that should not be discarded but could also lead to contradictions, if the shared peptides’ quantities do not fit to the quantities from the unique peptides. In the second example (N-shape, Fig 1B), protein B does not have a unique peptide and it is unclear, if it exists in the sample. Even if it does, it is not straightforward to quantify this protein, as it only has shared peptides whose quantities are also influenced by protein node A.

**Fig 1.**
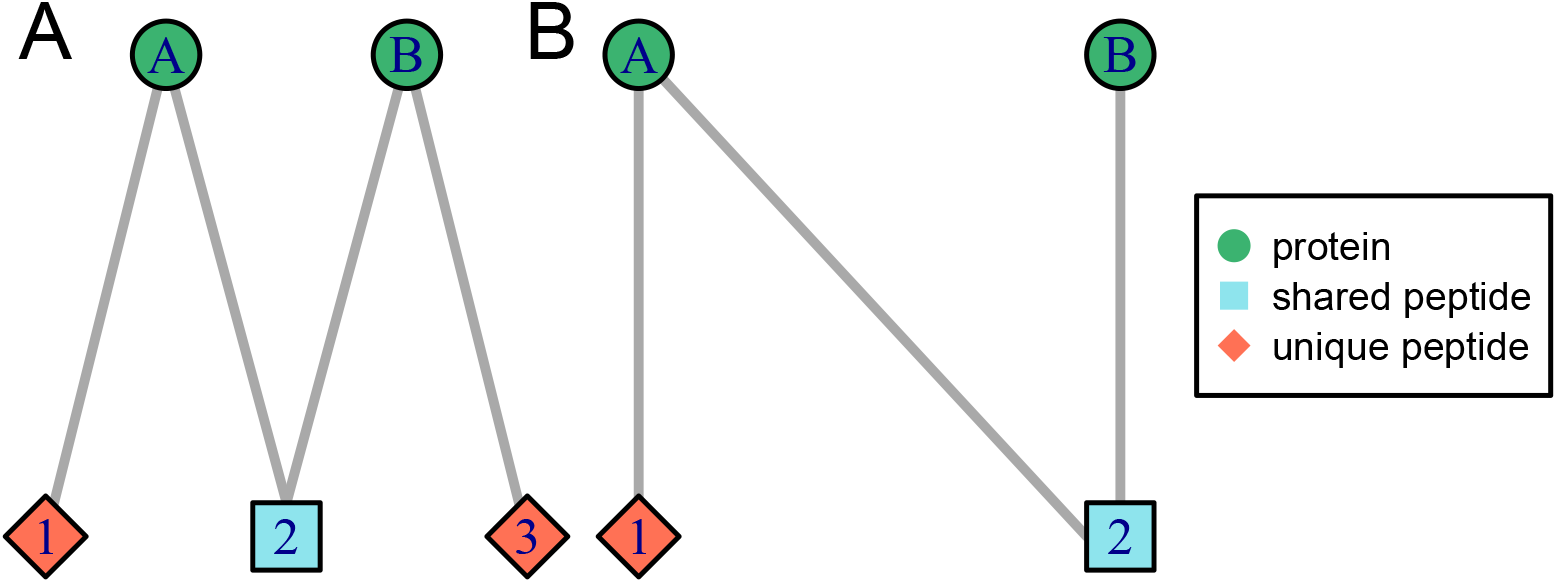
Examples of bipartite graphs depicting protein-peptide relationships. Protein nodes are shown as green circles and peptide nodes as red diamonds (unique to a protein node) or blue squares (shared between protein nodes). Each node may represent multiple protein accessions or peptide sequences. Unique peptide nodes are defined as nodes belonging to only one protein node, which may contain several protein accessions. Shared peptide nodes belong to at least two distinct protein nodes.

In this paper, we generate bipartite graphs of peptide-protein relationships using peptides quantified by mass spectrometry based proteomics in three different data sets as well as theoretical peptides from an *in silico* digestion of the related protein databases used for peptide identification. The aim of this paper is to systematically characterize the occurring bipartite peptide-protein graphs, to compare the theoretical graphs from the database with the ones from quantitative data sets. Furthermore, we can compare the different data sets and the databases from different species. We characterize common types of graphs as well as aggregated metrics over all graphs of a data set.

Bipartite graphs have been used in the past to display the relationships between proteins and peptides and to assist protein inference or quantification [12–15], a comprehensive analysis and characterization of the occurring graph types has yet not been conducted. Bamberger et al. [14] show the complete set of peptide-protein graphs of a combined data set from D. melanogaster and D. virilis. However, they do not characterize them further or compare them with theoretical graphs or different data sets, as it was not aim the of the paper (which is inferring proteoforms from the data).

This approach here serves as a model to describe the protein inference and quantification problem in more detail and assess its difficulty for a given data set. Especially the quantification of proteins without unique peptides is difficult or even not possible with most methods. We can show the extend of this problem with the help of the bipartite graphs, which differs between different data sets/organisms as well as between measured data sets and theoretical considerations of an *in silico* digestion of the corresponding databases. Results of this analysis will be helpful for development of new protein inference and quantification methods, especially for proteins without unique peptides.

## Materials and methods

### Data sets and corresponding protein databases

For building and analyzing the bipartite peptide-protein graphs, three different quantitative peptide-level data sets were used together with the corresponding protein databases. A summary of technical information and the search engine parameters for each data set can be found in S1 Table. The samples of the first data set (D1) consist of 13 non-mouse proteins that were spiked into a mouse cell background (C2C12 cell lysate) in different concentrations leading to five different states [16]. Each state was measured in three replicates leading to 15 samples in total. The raw files and the protein database were taken from the PRIDE repository [17], identifier PXD012986. Data were re-analyzed using a KNIME [18] workflow employing the search engines Mascot 2.7 [19], MS-GF+ [20] and X!Tandem [21]. Peptide identifications were combined using PIA [22, 23]. The workflow was similar to the original publication [16], except for a peptide identification filter directly after the spectrum identification leaving only peptides which are strictly tryptic (i.e. cutting after arginine (R) and lysine (K), if not followed by proline (P)). The resulting peptide quantities were used for further analysis (quantified_peptides-featureFinderCentroided.csv, which can be found in the PRIDE upload with the identifier PXD024684). The corresponding protein sequence database (in total 52,824 entries) consists of 52,548 entries from the UniProt reference mouse proteome (UP000000589, version 2017_12, only canonical sequences), the 13 spike-in proteins, 147 spike-in contaminants and 115 contaminants from the cRAP database.

For the second data set (D2) the UPS1 standard (containing 48 human proteins of equal molarity) were spiked into a yeast background in 10 different concentrations [24]. Raw files were taken from PRIDE, identifier PXD001819, and re-analyzed with MaxQuant version 1.6.17.0 [25, 26] and Andromeda [27]. Settings were taken from Workflow 7 of the corresponding publication [24], except that ”Trypsin” (no cutting after proline) was used instead of ”Trypsin/P”. The peptide output table from MaxQuant (peptides.txt file) was used for further processing of the quantitative peptide-level data. The corresponding database (in total 6,342 entries) consists of the UniProt yeast proteome (UP000002311, version 2019_11, 6,049 entries, only canonical sequences), the UPS1 fasta file [28] (48 entries), as well as the contaminants database [29] provided by Andromeda (245 entries).

Data set D3 is a re-analysis of a published ground-truth data set with two experimental groups. 60 *μ*g of HeLa cell lysate was mixed with either 10 *μ*g or 30 *μ*g of *E. coli* lysate, each measured in three technical replicates [26]. Raw files were taken from the PRIDE repository, identifier PXD000279, and re-analyzed with MaxQuant version 1.6.17.0 and Andromeda. The peptides.txt file was used for further processing of the quantitative peptide-level data. The database contains in total 81,709 canonical sequences and consists of the UniProt human reference proteome (UP000005640, version 2021_02, 77,027 entries), the *E. coli* reference proteome (UP000000625, version 2021_02, 4,437 entries) and the Andromeda contaminants database (245 entries). Additionally, we repeated this analysis including isoforms. The reference proteomes with isoforms contain 99,012 entries for human and 4,449 entries for *E. coli*. Together with the contaminant sequences, 103,706 sequences are included. This allows an analysis of the impact of including isoforms on the bipartite peptide-protein graphs.

For each of the protein databases, all protein sequences were digested *in silico* using the digestion rules for trypsin (cut after R and K, if not followed by P) to obtain all theoretical tryptic peptides. All peptides with a maximum of two missed cleavage sites were allowed. Only peptides with a length between five and 50 amino acids (AA) were considered further, as longer or shorter peptides are unlikely to be found via bottom-up MS [30]. Different values for the lower bound for the peptide length (five, six, seven and nine) were evaluated (see section *General overview and impact of minimal peptide length*).

### Processing of quantitative peptide-level data

For data set D1, peptides exclusively originating from decoy proteins were filtered out. Intensities for the same peptide sequence but with different post-translational modifications (PTMs) were summed up. Peptide sequences with a length larger than 50 AA or lower than the chosen minimal peptide length (see Section *General overview and impact of minimal peptide length* in the *Results* chapter) were removed, to be directly comparable with the theoretical peptides. The remaining sequences were mapped to the proteins in the corresponding database. The three replicates for each state were combined by averaging the peptide intensities, while zero values were treated as missing values (NA). If two or more of the three values were NA, the combined peptide intensity for this peptide was set to NA. For each possible comparison of the five different states, these aggregated values were used to calculate peptide ratios, which were set to NA if one of the corresponding values was NA. For each comparison, only peptides with a valid ratio (different from NA) were considered in the construction of the bipartite graphs (see Section *Construction of bipartite graphs*). For data sets D2 and D3, peptide LFQ values were extracted from the respective peptides.txt tables and zero values were re-assigned by NA. Further processing was the same as explained for data set D1, except that aggregation of peptides with different PTMs was not necessary here, as it is done internally by MaxQuant. An overview of the different quantitative peptide-level data sets can be found in S2 Table.

### Construction of bipartite graphs

For every data set, the bipartite protein-peptide graphs are constructed using two different sets of peptides. First, all theoretical peptides obtained from *in silico* digestion of the corresponding protein database are used (D1_fasta, D2_fasta, D3_fasta). Second, for each comparison of two states only those peptides with a valid ratio are used (D1_quant, D2_quant and D3_quant). The connected components that arise from the pairwise comparisons of the different states within one data set are put together and analyzed jointly.

To construct the bipartite graphs, a matrix with peptides in rows and all corresponding proteins in columns is created, with entries of 1 if the peptide belongs to the respective protein and 0 otherwise. This matrix is used as a biadjacency matrix [11] to build the bipartite graph of the relationship between proteins and peptides. Proteins that have the same set of peptides are indistinguishable and are therefore collapsed to a single node, i.e. a protein group. Similarly, all peptides sequences that belong to exactly the same set of proteins are collapsed to one single node to further simplify the graph structure. Accordingly, in Fig 1, one node may stand for multiple protein accessions or peptide sequences. This also means that here uniqueness is defined on the level of nodes. A peptide node (which may contain multiple peptide sequences) is defined as unique if it belongs to only one single protein node (which may contain multiple protein accessions), and it is defined as shared if it belongs to at least two protein nodes. In the resulting bipartite graph, protein nodes can be connected to each other via chains of shared peptide nodes. As not all protein nodes are connected this way, the graph can be decomposed into connected components, where all nodes are connected to each other, but not to any other node outside of it. The connected components (called ”graphs” in the following for simplicity) can be treated separately for protein inference or protein quantification.

Two graphs *G* and *H* are called isomorphic if there exists a bijective function *f* : *N*(*G*) → *N*(*H*) that maps the node set of *G* to the one of *H*, so that the edge structure remains the same: (*u*, *v*) ∈ *E*(*G*) ⇔ (*f* (*u*), *f* (*v*)) ∈ *E*(*H*) ∀ *u,v* ∈ *N*(*G*) [31]. Additionally, we require that the function *f* preserves the node type, i.e. that protein and peptide nodes are distinguished in terms of isomorphisms. Groups of isomorphic graphs reflect identical structures of protein-peptide relationships and contain the same information needed for inference or quantification, except the specific quantities of the peptides. Sets of isomorphic bipartite graphs (isomorphism classes) are built for the data sets, based on theoretical peptides from the *in silico* digestion of the database and quantified peptides with a valid ratio.

Classes of isomorphic graphs are determined by comparing two graphs using the ”isomorphic” function from the igraph package, which uses the BLISS algorithm proposed by Junttila and Kaski [32]. Both graphs are transformed into defined canonical forms which are then directly compared. These canonical forms are also used for graphical representation of the graphs in S2 Fig, S3 Fig, S4 Fig and S5 Fig. As the ”isomorphic” function does not take into account the type of nodes in a bipartite graph (e.g. a M-shaped and W-shaped graph would be considered isomorphic), we added a criterion that additionally compares the node types of both graphs. In an iterative procedure, each bipartite graph is compared to the current set of isomorphism classes and added to the corresponding class. If necessary, a new isomorphism class is built.

### Software

Calculations were performed and graphics created using R 4.1.2 [33]. The package seqinr 4.2-8 [34] was used to import the fasta files and OrgMassSpecR 0.5-3 [35] (modified version of the Digest() function) to perform the in *silico* digestion. For dealing with (bipartite) graphs, the igraph package 1.2.9 [36] was used, and for further graphics ggplot2 3.3.5 [37]. Additionally, function from the following R packages were used: reshape2 1.4.4 [38], BBmisc 1.11 [39], pbapply 1.5-0 [40], limma 3.47.16 [41], Matrix 1.3-4 [42], matrixStats 0.61.0 [43], openxlsx 4.2.4 [44], xtable 1.8-4 [45], tidyverse 1.3.1 [46], cowplot 1.1.1 [47], ggpubr 0.4.0 [48].

R code that was used for this work is available at: https://github.com/mpc-bioinformatics/bipartite-peptide-protein-graphs.

## Results

### General overview and impact of minimal peptide length

Very small and very large peptides cannot be measured by MS for high-throughput bottom-up proteomics [30]. During peptide identification, the length of peptides is usually restricted by limiting the number of amino acids as a setting for the search engine. Even if small peptides are allowed during database search, often only a few are identified and subsequently quantified in the end (see S1 Fig B, exemplary for D1). However, on database level, many small peptides exist after tryptic digestion. Especially peptides with a length of five amino acids (which are usually not identified nor quantified) show a larger proportion of shared peptides compared to longer peptides (see S1 Fig A, C and D). We analyzed the influence of different minimal peptide lengths on the overall bipartite graph structures (Table 1, S3 Table and S4 Table). We chose six and seven, as common default values for search engines, five as an extreme lower bound, and nine, the required length by the Human Proteome Project of the Human Proteome Organization (HUPO HPP) [49].

**Table 1.**
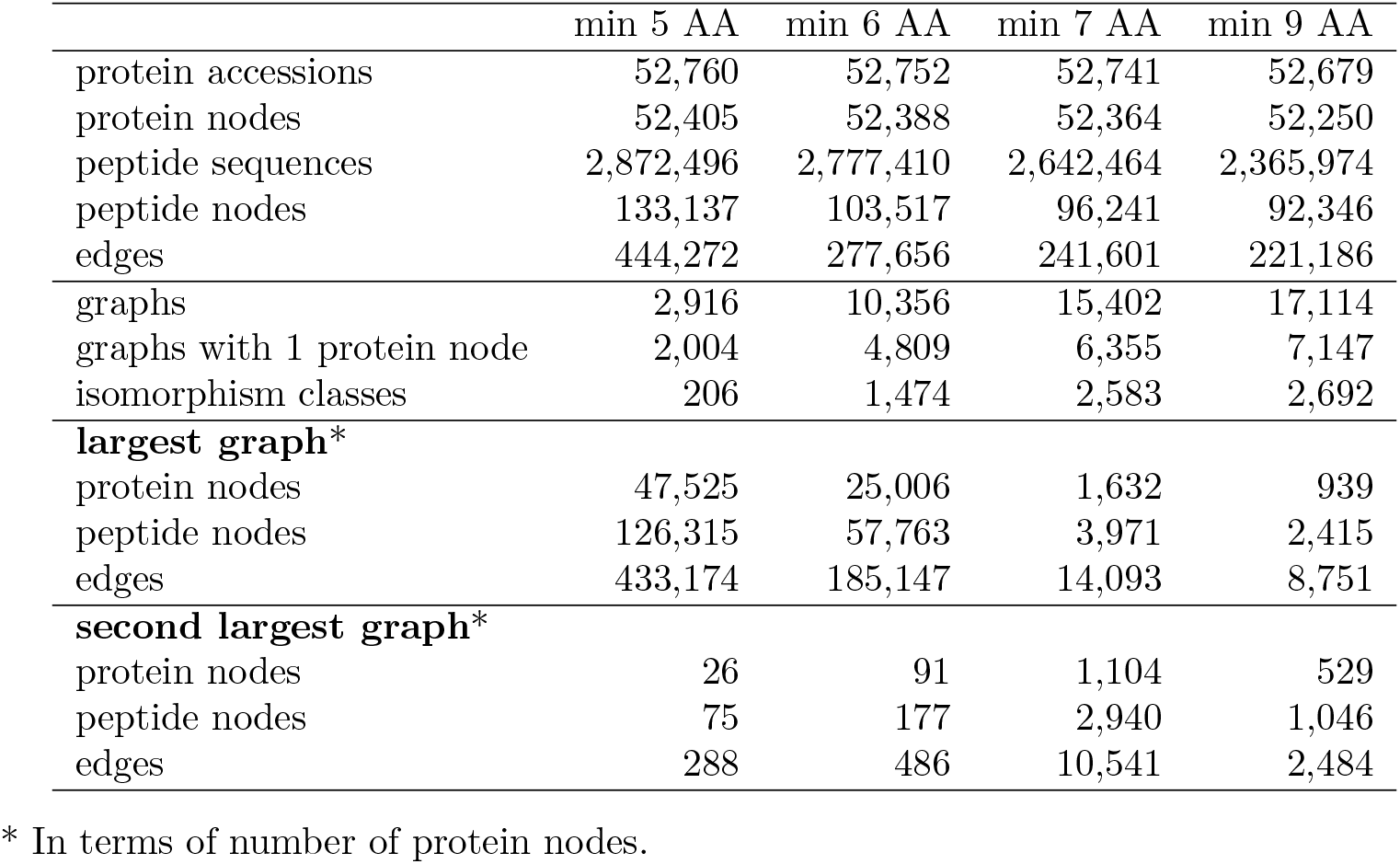
Influence of different minimal peptide lengths on the bipartite graphs for D1_fasta.

|  | min 5 AA | min 6 AA | min 7 AA | min 9 AA |
| --- | --- | --- | --- | --- |
| protein accessions | 52,760 | 52,752 | 52,741 | 52,679 |
| protein nodes | 52,405 | 52,388 | 52,364 | 52,250 |
| peptide sequences | 2,872,496 | 2,777,410 | 2,642,464 | 2,365,974 |
| peptide nodes | 133,137 | 103,517 | 96,241 | 92,346 |
| edges | 444,272 | 277,656 | 241,601 | 221,186 |
| graphs | 2,916 | 10,356 | 15,402 | 17,114 |
| graphs with 1 protein node | 2,004 | 4,809 | 6,355 | 7,147 |
| isomorphism classes | 206 | 1,474 | 2,583 | 2,692 |
| **largest graph\*** |  |  |  |  |
| protein nodes | 47,525 | 25,006 | 1,632 | 939 |
| peptide nodes | 126,315 | 57,763 | 3,971 | 2,415 |
| edges | 433,174 | 185,147 | 14,093 | 8,751 |
| **second largest graph\*** |  |  |  |  |
| protein nodes | 26 | 91 | 1,104 | 529 |
| peptide nodes | 75 | 177 | 2,940 | 1,046 |
| edges | 288 | 486 | 10,541 | 2,484 |

\* In terms of number of protein nodes.

For D1_fasta (Table 1), 52,824 protein sequences from the database were in *silico* digested to around 2.87 million peptides, including all peptides with a length between five and 50 amino acids and up to two missed cleavages. 64 protein sequences for which no tryptic peptide within the desired length was found were excluded from the analysis. The number of protein nodes is only slightly lower than the number of protein accessions (this is expected, as only few database proteins should be indistinguishable considering their tryptic peptides). However, the number of peptide nodes is notably lower than the number of peptide sequences (only about 130,000). The bipartite graph covering all these nodes consists of 2,916 connected components that form 206 different isomorphism classes. The largest class consists of the smallest possible graph with only one protein and one unique peptide node, which is an easy case for protein inference and quantification. This system occurs 2,004 times, making up around 69% of all graphs and covering 3.8% of all protein nodes. The largest graph (in terms of the number of protein nodes) contains over 47,000 protein nodes and therefore around 90% of all protein nodes and almost 95% of all peptide nodes. All these proteins are connected via chains of shared peptides. The second largest graph is much smaller with only 26 protein nodes and 75 peptide nodes.

Almost 100,000 short peptides of length five are included in the above analysis. When these are omitted (minimum peptide length six), the number of graphs more than triples to over 10,000. A major influence on this is the fragmentation of the former largest graph, which splits into over 7,000 graphs. Now, still about half of all protein and peptide nodes are connected within one single graph. The number of different isomorphism classes increases dramatically to almost 1,500. For peptides of length five the proportion of shared peptides is increased compared to longer peptides (S1 Fig), which can explain the breakdown of the largest graph. If the minimal peptide length is further increased to seven, the number of graphs rises further and the size of the largest graph decreases drastically from covering 47% of all protein nodes to only 3%. At this minimal peptide length, the largest graph reaches a reasonable size, also comparable to the second largest graph, without loosing too much information. Increasing the minimal peptide length even further, e.g. to nine, would lead to the loss of about 4,000 peptide nodes and 20,000 edges. We decided to use a minimal peptide length of seven, as a further increase risks losing too many well-quantifiable peptides and therefore complexity of the graph.

The second database (D2_fasta, S3 Table) contains 6,342 protein accessions, much less than D1_fasta. When using five AA as minimal length, only six protein accessions did not produce at least one peptide with the given criteria. More than 750,000 peptides are created during the *in silico* digestion. After collapsing of nodes, around 6,300 protein and 13,000 peptide nodes remain. 90.1 % of all graphs contain only one peptide node and protein node. There are only 18 different types of graphs. The largest graph contains 70.5% of all protein nodes and 84.1% of all peptide nodes. Increasing the minimal peptide length has a similar effect as for D1: the largest graph falls apart and the number of graphs rises. In contrast to D1, here the size of the largest graph already drops dramatically for a minimum peptide length of six (only contains 1% of all protein nodes). The number of protein nodes in the largest and second largest graphs are not influenced by omitting peptides of length seven and eight, however, they now contain fewer peptide nodes and edges.

For D3_fasta (S4 Table), based on human and *E. coli* databases, the behaviour is similar to D1_fasta, although on a higher level, because there are more protein accessions present in the FASTA file. Again, the number of graphs and of different isomorphism classes rises for higher minimum peptide lengths. As for D1_fasta, there is a large drop in size of the largest graph between six and seven amino acids (from 50% to 8% of all protein nodes). Between a limit of seven and nine, again the increase has slowed down compared to lower limits.

In summary, the minimal peptide length has a huge impact on the graph structures and especially on the size of the largest graph. For the following analyses, a minimal peptide length of seven for D1_fasta and D3_fasta and of six amino acids for D2_fasta is used. This ensures that the largest graph is decomposed far enough to yield a reasonable size compared to the total number of protein nodes. On the other hand we do not omit too many peptides and therefore retain valuable information. These thresholds are also applied to the quantitative peptide-level data to ensure direct comparability.

### Occurring types of graphs and aggregated characteristics

In the following, common isomorphism classes occurring from theoretical or quantified peptides of D1, D2 and D3 are described. Aggregated characteristics of all graphs are calculated as a summarization and shown in S2 Fig and Fig 3. For D1_fasta, 15,402 bipartite graphs are constructed which fall into 2,583 different isomorphism classes. The representative graphs of the ten largest isomorphism classes are shown in Fig 2A. The quantitative data set D1_quant contains five states of different combinations of concentrations of 13 spike-in proteins, leading to ten possible pairwise comparisons. Over the ten comparisons 28,802 graphs are formed in total, which belong to 354 different isomorphism classes. To characterize the graphs that occur, the distribution of numbers of different node types is shown in S2 Fig (first row). The two main node types are protein and peptide nodes, which can be further divided into unique and shared peptide nodes. The most simple graph type is the only one with just one protein node. This is the most frequent class for D1_fasta and D1_quant with 41% and 61% of the graphs, respectively, and therefore the corresponding bar is the highest. Moving from the database to the quantitative level, the proportion of this simple graph type increases due to the disappearance of shared peptides that are not quantified (see Section *Processing of quantitative peptide-level data*). While this graph type contains 12.1% of all protein nodes in the theoretical setting, this proportion rises to 32.5% for D1_quant.

**Fig 2.**
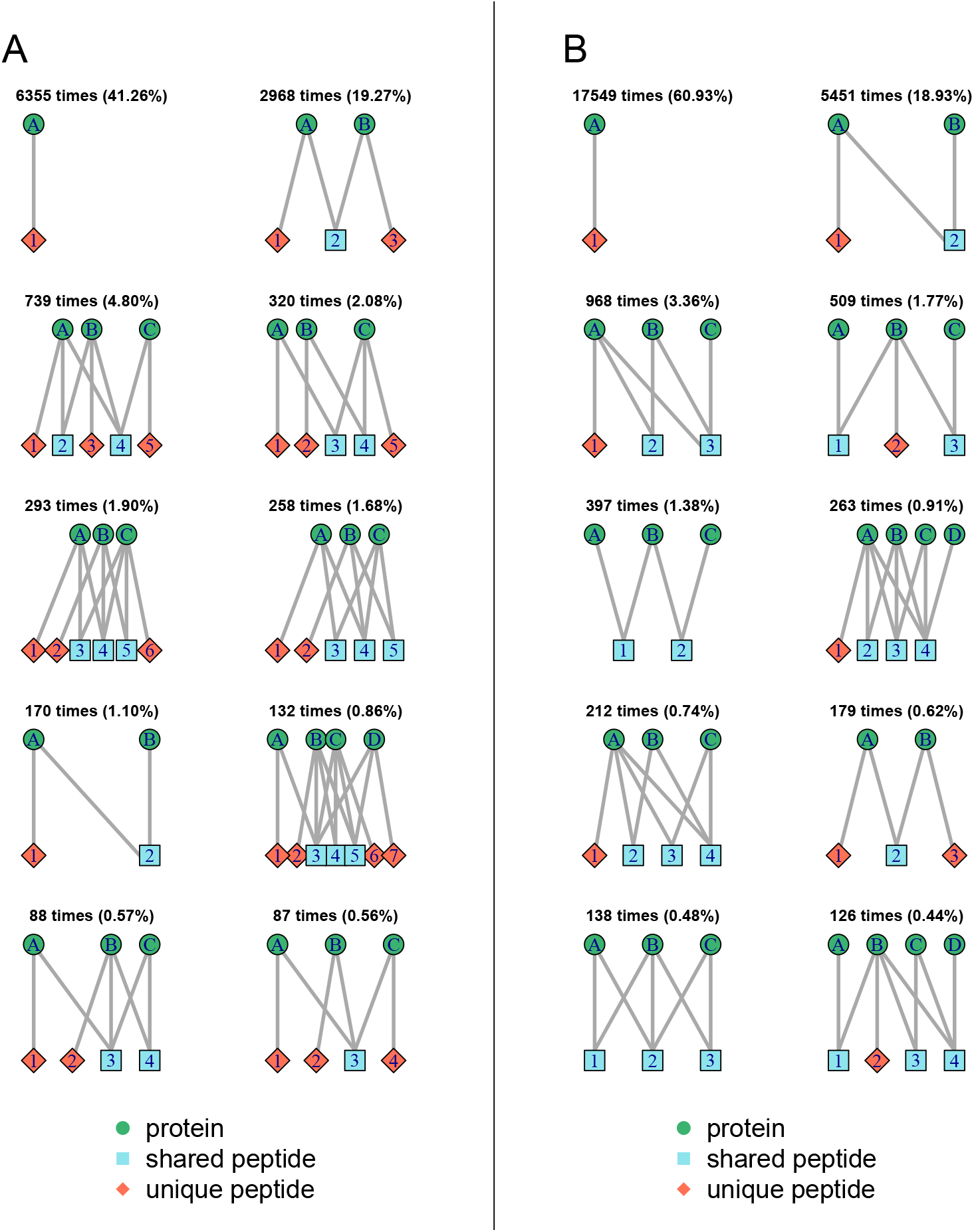
Representative bipartite graphs for the ten largest isomorphism classes found in data set D1. A) D1_fasta, B) D1_quant, with number of occurrences and percentage of all graphs.

**Fig 3.**
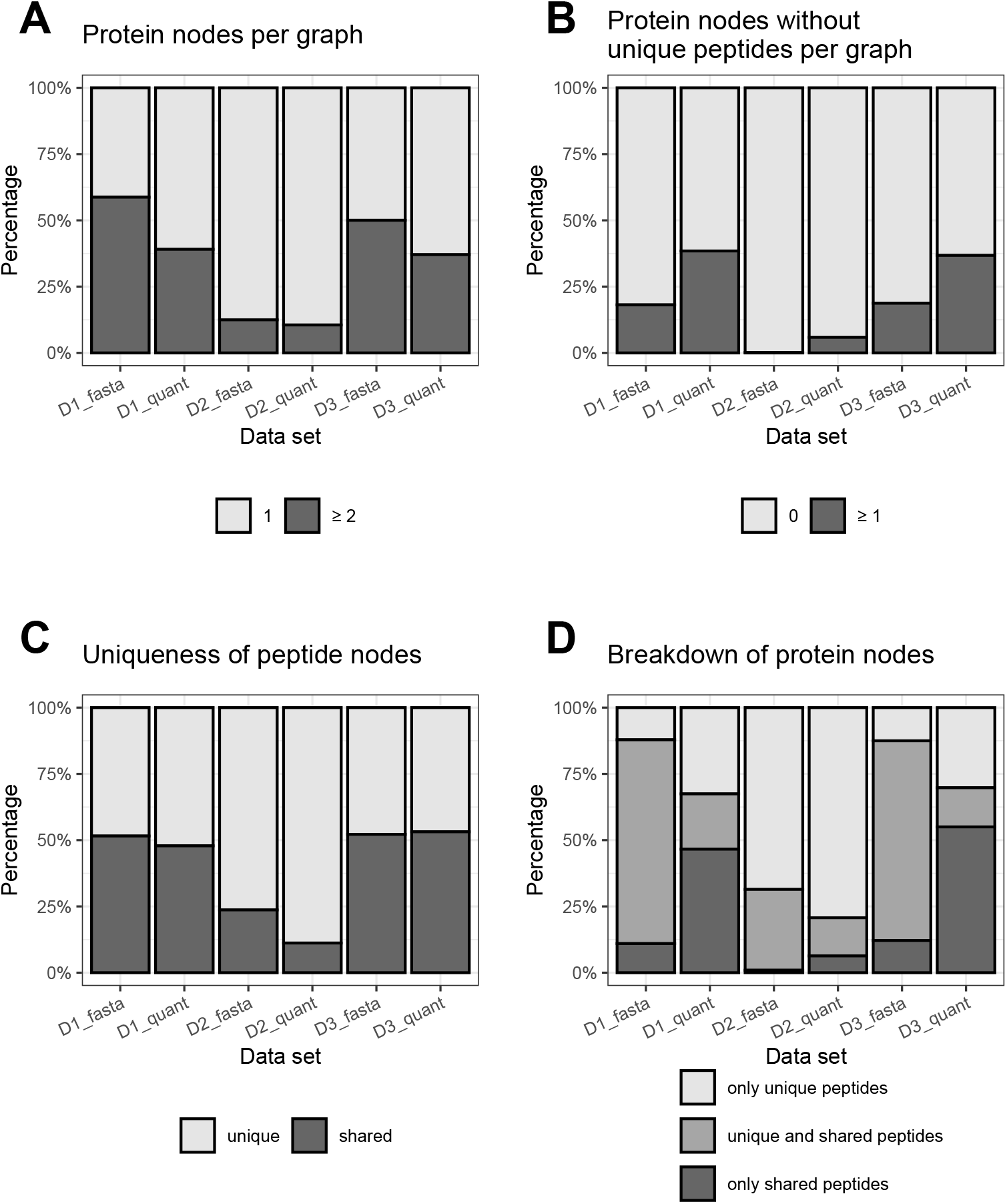
Aggregated characteristics of bipartite peptide-protein graphs over the different data sets. A: Percentages of graphs with one or with more than one protein node. The light grey bars reflects the percentage of the most simple graph type with only one protein node compared to all graphs. The dark grey bars represents the percentage of all larger graphs with two or more protein nodes. B: Percentages of graphs with none vs. one or more protein nodes, that are not connected to any unique peptide node. In the graphs represented by the light grey bars, each protein node has at least one unique peptide. In the graphs represented by the dark grey bars, at least one protein node exists without a unique peptide (which is a more complicated case for protein inference and quantification). C: Percentages of unique vs. shared peptide nodes in relation to all peptide nodes across all bipartite peptide-protein graphs. Uniqueness is here defined as belonging to only one protein node, which may consist of multiple protein accessions. D: Percentages of protein nodes with only unique peptides vs. unique and shared peptides vs. only shared peptides in relation to all protein nodes across all bipartite peptide-protein graphs. Uniqueness of the peptide nodes is here defined as belonging to only one protein node, which may consist of multiple protein accessions. Please note: As described in the ‘Methods’ section, peptide nodes may contain multiple peptide sequences and protein nodes may contain multiple protein accessions.

The second largest class for D1_fasta (almost 3,000 occurrences, i.e., around 19% of the graphs) contains graphs with two protein nodes and three peptide nodes. In these M-shapes graphs there are two peptide nodes that are unique for one of the protein groups each, and a peptide node that is shared by the two protein nodes. As each protein group has at least one unique peptide, both should be reported during protein inference. For quantification, however, it is not clear how to handle shared peptides. They may be omitted (as done in some currently used quantification algorithms), but this would mean to delete valuable information. While this graph type is common in D1_fasta, its proportion strongly decreases for D1_quant to only 0.62%. Among the ten largest isomorphism classes for D1_fasta, there are four (ranks three, four, five and ten) that have three protein nodes that are each connected to a unique peptide node. The graphs differ in number of and connections to shared peptide nodes. From an inference and quantification point of view, they behave similar to M-shaped graphs. The same holds for the eighth most common isomorphism class, which contains four proteins, each connected to a unique peptide node. The sixth, seventh and ninth largest isomorphism classes have in common, that one of the protein nodes is not connected to a unique peptide node. This means that from the identified peptides it is not clear if this protein is in the sample or not. However, the quantities of the shared peptides in combination with the unique peptides can help to estimate a quantity for this protein node. E.g., in case of N-shaped graphs (see Fig 1B) with only two protein nodes and one unique and one shared peptide node, this is possible. The ratio for protein B cannot be given exactly, but a lower or upper limit can be calculated, depending on the values of the two peptide ratios. This opportunity is not exploited by most currently applied protein quantification methods. Of the remaining isomorphism classes, 2,306 only occur once in D1_fasta. These are often relatively large systems that are uniquely found in the data due to their complex combination of protein and peptide nodes and the corresponding edges. However, these graphs contain almost 26,700 protein nodes which corresponds to almost 51% of all protein nodes and should not be neglected. For the quantitative data set D1_quant graphs with a unique peptide node for each protein node are not within the ten largest classes (except the M-shaped graph) and occur only seldomly. Instead, from the top ten graph types, eight contain at least one protein node without a unique peptide node. The N-shaped graph is the second most common with almost 19% of all graphs The fifth and ninth largest isomorphism classes do not contain any unique peptide node. These differences between the theoretical graphs from D1_fasta and the graphs from the quantitative data set D1_quant can also be seen in S2 Fig.

On database level, graph types with two peptide nodes (N-shaped, like Fig 1B) are very uncommon and much less present than those with three peptide nodes. This graph type forms the second largest class for D1_quant, but almost never appears on the database level. The M-shaped graph type (see Fig 1A) is the second most common on the database level, which causes a higher bar for a peptide number of three in S2 Fig. A lot of M-shaped graph types fall apart from database to quantitative level to the most simple or N-shaped graphs or vanish completely. In general, for the database level, graph types with even numbers of peptide nodes seem to be less common than the following odd numbers. This causes ”gaps” for even numbers in the peptide node distributions for theoretical peptides, while these are not present for quantified peptides. For unique or shared peptide nodes these gaps do not exist. In this case, also the number zero is possible for the smallest graph type. Zero unique peptides occur rarely, but a little more often on the quantitative than on the database level. These graphs are complex during protein quantification, as there is no unique peptide as an ”anchor”.

For D2_fasta, 4,908 graphs are formed that fall into 64 different isomorphism classes. Again, the most common graph type is the most simple one (S3 Fig (a)). It is more common than in D1_fasta with more than 87.5% of all graphs that cover in total 68.6% of all protein nodes. This can be explained by the lower complexity of the analyzed yeast compared to the mouse proteome, which leads to fewer connections between proteins via shared peptides. The second largest class also is the same as for in D1_fasta (M-shaped graph), with 8.54% of all graphs. The remaining classes are each relatively seldom but nevertheless contain around 18% of all protein nodes. In all of the top 10 graph types all proteins nodes are connected to a unique peptide node.

On quantitative level (D2_quant, S3 Fig (b)), the percentage of the most simple graph type only increases slightly compared to the database level. This increase is less pronounced compared to data set D1, since the proportion of the simple graph type is already very high on database level for D2_fasta, see also Fig 2. However, the N-shaped graph type (which did not exist at all in D2_fasta) is now the second most common, like for D1_quant, with 4.77%. The M-shaped graph type moves from place two to three and still covers 4.56% of the graph types. Seven of the graph types in the top ten now contain proteins without any unique peptide, which was not seen on database level. In general, the decreasing frequency of larger graphs is the strongest for D2 among the compared data sets, especially for the quantitative level, where the distribution of the number of nodes is shifted to smaller numbers, as graphs split up due to disappearing shared peptides (see Fig 2).

The graphs for data set D3 (S4 Fig (a) and (b)) behave more similarly to D1 than to D2. While the percentage of the most simple graph type is higher for D3_fasta compared to D1_fasta (almost 50% compared to 41%), this value is quite similar for D3_quant and D1_quant (63% and 61%, respectively). Overall, the graph structure and even the order of the ten largest isomorphism classes is nearly the same for D3_fasta and D1_fasta. On the quantitative level it can be noted that nine of the top ten classes contain protein nodes without unique peptides and there are three with only shared peptides. The M-shaped graph is missing from the top ten here. The patterns shown in Fig 2 are also very similar between D1 and D3. Larger graphs with ten or more nodes occur more often in D3. Graphs without any unique peptide also occur more often in D3_quant than in any other of the analyzed quantitative data sets.

A summarization of these findings can be found in Fig 3. In general, the percentage of the smallest graph type (Fig 3A, light grey bar) increases when going from database to quantitative level. For D2, the difference is rather small, as the proportion of the smallest graphs is already high on database level (87.5% vs. 89.5%). For D1 and D3, this proportion increases much stronger, leaving a large number of systems (with an even larger number of affected protein nodes) that are influenced by shared peptides. Protein nodes without a unique peptide node are the most difficult to quantify and are ignored completely or subsumed into protein subgroups by many quantification methods. The percentage of graphs that contain at least one of these protein nodes (3B, dark grey bar) increases on the quantitative level in comparison to the database level, possibly explained by the disappearance of non-quantified unique peptides. For D2, the increase is on a much lower level compared to D1. For D3, the proportion of graphs that contain at least one protein node without unique peptides is the highest.

Fig 3C and D show the distribution of uniqueness for peptide and protein nodes. The proportion of shared peptides slightly decreases from database to quantitative level and is by far the lowest in D2. While the proportion of protein nodes with unique and shared peptides both decrease from database to quantitative level, the proportion of nodes with only one of the two peptide node types increases. Especially for D1 and D3, the frequency of protein nodes with only shared peptides strongly increases. These proteins are hard to quantify, as they are missing unique peptides and the shared peptides are also influenced by other proteins. On the quantitative level, 46.6% and 55.0% of the protein nodes fall into this category for D1_quant and D3_quant, respectively, while this proportion is only 6.3% for D2_quant.

### Influence of the inclusion of isoforms

For data set D3 we conducted the analysis using two different protein sequence databases. As described in section *Data sets and corresponding protein databases*, the reference proteome databases from human and *E. coli* were downloaded from UniProt once containing only canonical sequences and once also including isoforms. In total, the database with isoforms contains almost 22,000 additional entries (12 from *E. coli*, the remainder from the human part of the database). We chose D3 for this analysis because the human database contains the most annotated isoforms among the three analyzed species (for mouse, around 7,500 and for yeast 29 isoforms are annotated for the corresponding reference proteomes). We expect the largest effect of adding isoforms for this data set.

A summary of the characteristics of the generated bipartite peptide-protein graphs with and without isoforms, on database and quantitative level, can be found in S5 Table. The *in silico* digest of the database with isoforms now contains 27% more accessions, only about 100,000 new peptides are generated, i.e. 3.4%. It can be observed that over 500,000 of formerly unique peptides are shared after adding isoforms, but the large majority remains isoform-unique, i.e. peptides are shared but only between a canonical sequence and its isoforms (see Table 2). From the new peptide sequences, the majority – around 90% – is unique for one isoform. In total there is a shift from 43.2% shared peptides in the canonical database to 58.4% in the database including isoforms (also including isoform-unique peptides). It can be observed that the number of bipartite peptide-protein graphs slightly decreases when isoforms are added. Two formerly separate graphs may be connected by an isoform if it shares peptides with proteins from both graphs. The number of graphs with only one protein node decreases by 11.6%.

**Table 2.** Unique and shared peptides stemming from *in silico* digestion of D3_f’asta and D3_iso_fasta (without and with isoforms).

| with isoforms | without isoforms |  |  | sum |
| --- | --- | --- | --- | --- |
|  | unique | shared | not existing |  |
| unique* | 1,218,900 | 0 | 93,856 | 1,312,756 |
| isoform-unique** | 485,723 | 0 | 10,087 | 495,810 |
| shared | 26,027 | 1,319,690 | 103 | 1,345,820 |
| sum | 1,730,650 | 1,319,690 | 104,046 | 3,154,386 |

\* Unlike the rest of this paper, unique is defined here as unique for a protein accession. However, the difference between protein nodes and accessions is negligible when looking at the database level.
\*\* Isoform-unique peptides are defined as shared peptides that are shared only between isoforms of the same protein.

Allowing isoforms in the database leads to a slight decrease of quantified peptides. Again, the number of graphs decreases. The largest ten isomorphism classes do not change heavily, but it can be observed that graph types without any unique peptide become more frequent (e.g. the W-shaped graph that rises from 2.73% to 3.80%, see S5 Fig (a) and (b)). As a conclusion, while the number of graphs decreases slightly, the graphs become slightly larger and there are more protein nodes without unique peptides when including isoforms (see Fig 4).

**Fig 4.**
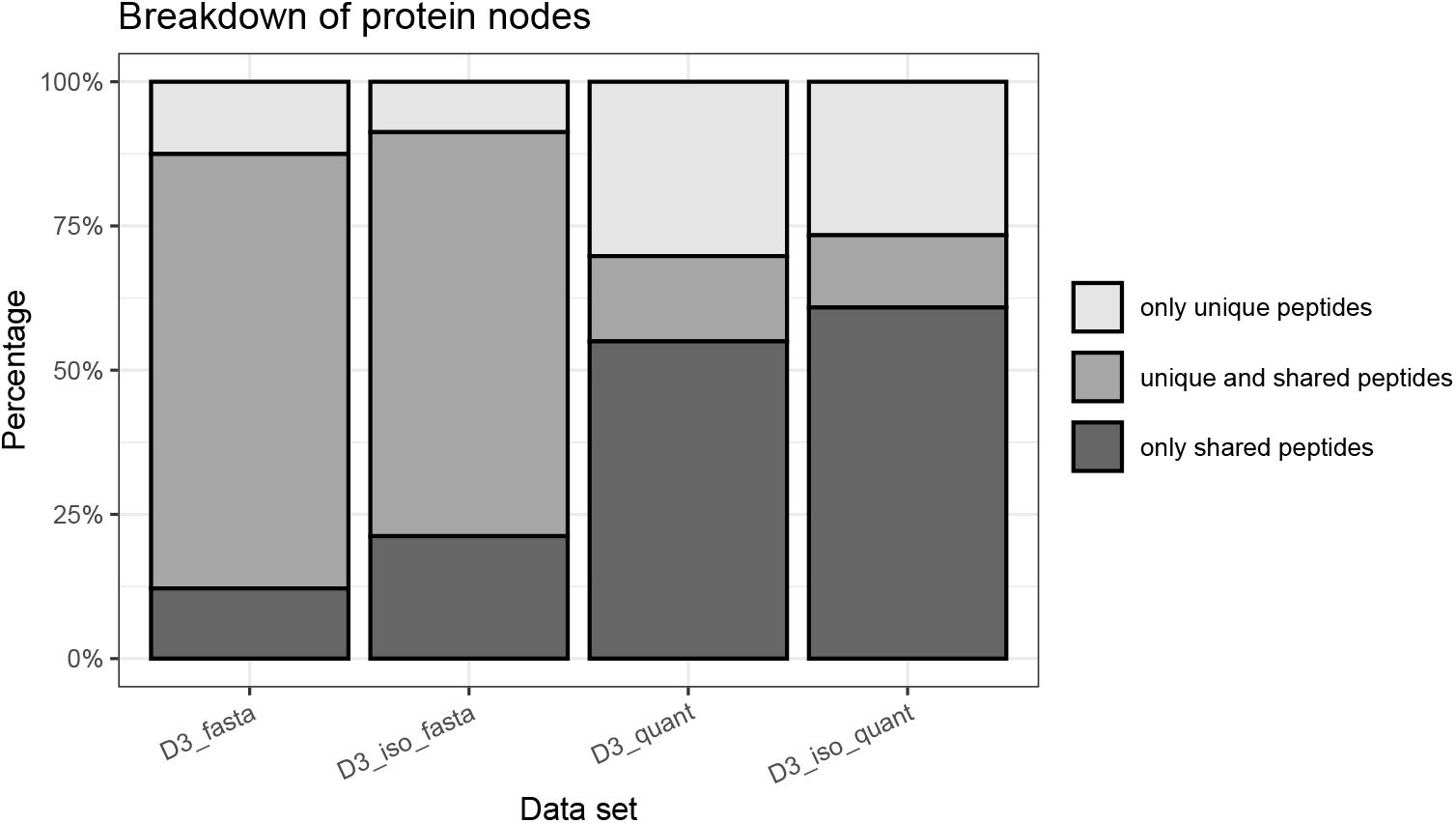
Distribution of protein node types with and without the inclusion of isoforms. Percentages of protein nodes with only unique peptides vs. unique and shared peptides vs. only shared peptides for data set D3, with and without isoforms. Please note: As described in the ‘Methods’ section, peptide nodes may contain multiple peptide sequences and protein nodes may contain multiple protein accessions.

## Discussion

Despite the large number of methods and tools available for protein inference and quantification, there is currently no commonly accepted method for dealing with shared peptides. Especially the problem of quantifying proteins with only quantified shared peptides is often overlooked by subsuming the protein into a protein group together with the other proteins of the shared peptide. Bipartite peptide-protein graphs have been used in the past to illustrate the protein inference and quantification problem. Here, we used the characterization of these graphs as an approach to assess the difficulty of this problem for a given data set. The investigation of bipartite peptide-protein graphs is therefore highly relevant to the current research in proteomics. Furthermore, the analysis of bipartite graphs can also be applied to the field of proteoform research by including isoforms and peptides with PTMs [50].

In our study we saw that the minimum peptide length is a crucial setting for the *in silico* digestion of proteins and has a huge impact on the structures of the bipartite graphs. Allowing too small peptides leads to large graphs that cover a majority of the proteins inside a database which is not informative. It also does not reflect the situation for real measurements, as small peptides are rarely found or even searched for in database searches. The minimal peptide length that fits best to the data also varies by database and organism. Here, we have chosen the smallest peptide length which caused a large drop in the size of the largest graph and leads to a manageable size of it (in the analyzed data covering between 1 and 8% of all protein nodes). We selected a minimal peptide length of seven for D1 and D3 and six for D2 for further investigation. Smaller peptides were not or very rarely quantified in the data sets and lead to a largest graph that covers the majority of protein nodes. If such a large graph would really occur on the quantitative level, this would be a complicated case for protein quantification, because of the high complexity of the interactions of the different proteins via small shared peptides. If we further increase the minimal peptide length on the other hand, we would omit a high number of edges and lose important information about the connection between proteins via shared peptides, that we want to keep. Although we used a smaller threshold for D2 (six amino acids), the impact of increasing it to seven amino acids is very small and neglectable, so we conclude that we can still compare the findings with those from D1 and D3. In favor of using more peptides that connect proteins in a data set with comparably few shared peptides, we decided to continue with a threshold of six rather than increasing it also to seven for D2. The qualitative message with a large difference between D2 compared to D1/D3 is not influenced by this decision.

To reflect the situation within a peptide search engine, we *in silico* digested the whole protein database that was used for searching the respective raw data. We decided to include all possible tryptic peptides within the desired length range. This also includes all possible peptides with up to two missed cleavages, if they do not exceed the upper limit of 50 amino acids. As a result, the number of peptides is blown up. However, it is exactly the situation that the search engines face. Often, the miscleaved peptides behave similar to the smaller counterparts in regard to uniqueness and will therefore often be contained in the same peptide node. However, we still capture the case in which a longer miscleaved peptides is unique while the shorter version is not.

In all analyzed settings, the most simple graph type with only one protein and one peptide node is the most frequent one. In this case, the sequences in the peptide node are unique to the protein node, which may consist of multiple protein accessions that could not be distinguished by their peptides (i.e. they form a protein group). These systems are an easy case for protein inference or quantification. As there are unique peptides, there is strong evidence for the presence of the corresponding protein (or at least one of the proteins within the node). A simple strategy for quantification, like averaging the peptide quantities, may work well in these cases. However, one has to keep in mind that no matter which quantification method is used, this will only lead to a common quantity for all the members of a protein node if it consists of multiple protein accessions.

The composition of graph types differs between database and quantitative level. Larger bipartite graphs split up into smaller ones when going from the database to the quantitative level, while at the same time graph types containing proteins without any unique peptide become more prominent. Aggregated characteristics over the whole set of bipartite graphs confirm these findings. There are two opposing effects when comparing database level and quantitative level. First, the proportion of the smallest possible graph (situation with simple solution for quantification) becomes larger. Second, the proportion of proteins without unique peptides is considerably higher on quantitative than on the database level. For these proteins, inference and quantification becomes more challenging, especially in cases where there is no unique peptide in the graph at all. This indicates the importance of considering the quantified peptides instead of assessing the difficulty of quantification only based on the corresponding databases. The reason for the huge differences between database and quantitative level is that many theoretical peptides remain unquantified. These missing peptides cause the graphs to fall apart if these are shared peptides. Missing unique peptides cause some protein nodes to completely lose their unique peptides. These protein nodes may collapse with another node with the exactly same set of quantified peptides, thus leading to a smaller graph. In other cases, they remain in a protein node without unique peptides, which is a challenge for protein quantification algorithms.

We defined the uniqueness of peptides on the level of nodes, i.e. a peptide is considered unique if it belongs to only one single protein node. Especially on the quantitative level, this protein node may contain multiple protein sequences/accessions that are indistinguishable regarding their quantified peptides. We have chosen this definition because protein accessions within one node are not distinguishable regarding their (quantified) peptides and therefore in any case only one quantity for this protein group can be calculated.

There are also large differences in the characteristics of the bipartite graphs between the three different data sets for theoretical as well as quantified peptides. For example, the analysis of D2 leads to a higher proportion of unique peptide nodes and a more frequent occurrence of the most simple graph type. This makes the overall protein inference and quantification considerably easier than for D1 or D3. A possible explanation could be the different complexities of the predominantly underlying organisms, mouse (D1), yeast (D2), and human (D3). The yeast genome is clearly smaller than the mouse or human genome (around 12 Mb [51] vs. over 2,600 Mb [52] vs. over 2,800 Mb [53], respectively), and the same holds for the proteome (around 6,000 vs. over 50,000 vs. over 75,000 protein entries without isoforms in the UniProt reference proteomes, respectively). Already on the database level, the percentage of unique peptide nodes and of protein nodes with only unique peptides for D2 is much higher than for D1 or D3. This indicates that the complexity of protein quantification can vary a lot for different organisms and data sets, which may also have an impact on the choice of a suitable protein quantification method. For D2 a simple quantification method, like averaging the unique peptides, might work well for most proteins. For D1 or D3, in contrast, this would leave many proteins unquantifiable (46.6% and 55.0%, respectively) because of the lack of unique peptides and would also ignore a lot of information from the large number of shared peptides (47.9% and 53.2%, respectively) (see Fig 3C and D). In this case a more appropriate method that takes into account the shared peptides and the graph structures is crucial to get the most out of the acquired peptide data. We showed that the underlying organism has a huge effect both on the database and the quantitative level. As a consequence, this kind of study should be repeated for other organisms of interest. This again highlights the need to generate the bipartite graphs also for only the quantified peptides and not only for all possible peptides from the database, and then to compare these two scenarios.

One has to keep in mind, that not all three data sets were measured with the same type of mass spectrometer. While data sets D2 and D3 were measured with an LTQ-Orbitrap instrument, D1 was measured by a Q Exactive HF, both by Thermo Fisher Scientific. Of course, using a different instrument type can have a large impact on the number and accuracy of the quantified peptides. However, we see large differences between the three organisms already on the level of theoretical graphs using the FASTA files, without any quantitative information. Furthermore, also on the graphs from the quantitative data, D1 and D3 behave very similar in our analysis, even though they are measured on different instrument types. There are large differences between D2 and D3, although they were measured on a comparable machine. Therefore, we can conclude that the largest differences are indeed due to the different complexities of the organisms (yeast vs. human). However, a comparison of the same data set measured by different mass spectrometer types is necessary to fully elucidate the impact on the bipartite graphs.

All three data sets were measured in the data-dependent acquisition mode (DDA), so they are comparable in this regard. We expect differences for data from data-independent acquisition (DIA), because usually here more peptides are identified and quantified [16]. This may lead to more unique peptide nodes, which make the inference and quantification of the proteins easier. On the other hand, more shared peptides may be quantified, which may connect graphs to each other. For spectral library-based DIA it would also make sense to compare the graphs from the quantitative data with the ones generated by the peptides in the spectral library, in addition to the whole FASTA database.

One has to be careful though, as different specifications (e.g., enzyme for digestion, mass spectrometer type, fragmentation and ionization techniques) may have an impact on the resulting graphs on the quantitative level, although this was not investigated here. Additionally, on the database level the complexity may appear higher or lower dependent on small variations of the settings (e.g., minimal peptide length, allowed number of missed cleavages). The additional consideration of isoforms was performed on data set D3 as an example. While this makes the graphs more complicated and rises the number of protein nodes without unique peptides, isoforms may explain otherwise dubious peptide intensities and can be of high interest for the biological research question. We did not investigate the influence of peptides with PTMs here, but the bipartite graphs could be helpful for assessing the situation for quantifying different proteoforms with the same peptide sequence.

Independent from the peptide quantification method applied, the characteristics shown in this study can easily be generated and interpreted. In the future, different MS techniques might turn out to generate graphs more close to the database level than others, which may have a huge impact on the quality and difficulty of protein quantification. To reach the same graph shape as on the database level, not all of the theoretically possible peptide sequences need to be identified and quantified but at least one peptide sequence per peptide node, which is much more realistic.

Besides protein and peptide nodes there may also be a third type of nodes, the MS2 spectrum nodes. MS2 spectra can be chimeric, i.e. containing the fragment ions of two or more co-eluting precursors. In a 3-partite graph, these nodes may therefore build new connections between peptide nodes. In this manuscript, we ignored these potential connections (in about 1% of the spectra a second peptide was identified). Depending on the measurement technique and algorithms used, it may be possible or not to separate the MS1 level quantifications for different precursors inside the same MS2 spectrum. It should be further investigated what implications these connections inside the 3-partite graphs may have for protein quantification.

## Conclusion

In this paper, we analyzed bipartite graphs of protein-peptide relationships using three different quantitative data sets, originating from different(ly complex) species, as well as the corresponding protein databases. Our aim was to better understand the relationship between proteins and peptides and the implications for protein inference and quantification. In general, looking at bipartite peptide-protein graphs and aggregated characteristics is helpful to judge how complex the protein inference and quantification will be for a particular data set. It can also give a hint whether simple protein quantification methods like averaging the unique peptide intensities would lead to a sufficient proportion of quantified proteins or if they would miss a large potential of the data. Depending on the data set, shared peptides can make up a large proportion of all quantified peptides, especially when isoforms are considered. This is accompanied by a large number of protein nodes without unique peptides, which are often overlooked by protein quantification algorithms. As a consequence, there is an urgent need for protein quantification methods that take into account the graph structures and that can use this information to reliably quantify proteins without unique peptides. In conclusion we recognize three main benefits from our study: 1) the systematic characterization of occurring bipartite peptide-protein graphs, which has not been done before, 2) the gain of insights about the impact of the choice of the minimal peptide length for the *in silico* digestion and 3) the recognition of the potential for using these insights for development of protein inference and quantification algorithms.

## Acknowledgments

The authors thank Anika Frericks-Zipper for help with the PRIDE upload.

## Availability of data and materials

The datasets supporting the conclusions of this article are available in the ProteomXChange Consortium at http://proteomecentral.proteomexchange.org via the PRIDE partner repository and can be accessed with the dataset identifiers PXD024684 (D1), PXD030577 (D2) and PXD030603 (D3).

## Authors’ contributions

K.S.: Conceptualization, Methodology, Software, Formal Analysis, Writing – Original Draft, Visualization

M.T.: Conceptualization, Methodology, Validation, Writing – Review & Editing

J.U.: Formal Analysis, Writing – Review & Editing

J.R.: Conceptualization, Methodology, Writing – Review & Editing, Supervision

M.E.: Conceptualization, Methodology, Writing – Review & Editing, Supervision, Funding acquisition

All authors read and approved the final manuscript.

## Supporting information

**S1 Table. Technical information and search engine parameters for the three analyzed quantitative proteomics data sets.**

**S2 Table. Overview over the quantitative peptide-level data sets.**

**S3 Table. Influence of different minimal peptide lengths on the bipartite graphs for D2_fasta.**

**S4 Table. Influence of different minimal peptide lengths on the bipartite graphs for D3_fasta (without isoforms).**

**S5 Table. Comparison of bipartite graph characteristics without and with isoforms on data set D3 (with minimal peptide length of seven amino acids).**

**S1 Fig. Count and percentages of shared and unique peptide sequences depending on the peptide length.** The peptide length is given in amino acids. As an example, here the values for data set D1 are shown. Uniqueness is here defined as belonging to only one protein node, which may consist of multiple protein accessions.

**S2 Fig. Distribution of numbers of different node types for the bipartite peptide-protein graphs.** The leftmost column shows the distribution of number of protein nodes for D1, D2 and D3, on database level (*_fasta) and on the level of quantified peptides (*_quant). On the x-axis the number of protein nodes are shown and on the y-axis the proportion of graphs with this exact number of protein nodes in comparison to all graphs. E.g., the bar at x = 1 is the proportion of graphs with only one protein node compared to all graphs. The sum of all bar heights adds up to one for each subfigure. Similarly, the distribution of the number of total peptide nodes, unique peptide nodes and shared peptide nodes are shown in the following columns. Minimal peptide lengths were chosen as seven for D1 and D3 and six for D2. The rightmost bar with the label ”10+” comprises the values of 10 and above.

**S3 Fig. Representative bipartite graphs of the ten largest isomorphism classes found in data set D2.** (a) D2_fasta, (b) D2_quant, with number of occurrences and percentage of all graphs.

**S4 Fig. Representative bipartite graphs of the ten largest isomorphism classes found in data set D3** (a) D3_fasta, (b) D3_quant, with number of occurrences and percentage of all graphs.

**S5 Fig. Representative bipartite graphs of the ten largest isomorphism classes found in data set D3_iso.** (a) D3_iso_fasta, (b) D3_iso_quant, with number of occurrences and percentage of all graphs.

