## Supplement for "Characterization of peptide-protein relationships in protein ambiguity groups via bipartite graphs"

### Supplementary Tables and Figures

**S1 Table:** **T**echnical information and search engine parameters for the three analyzed data quantitative proteomics data sets.

| Parameter | D1 | D2 | D3 |
|---|---|---|---|
| Mass spectrometer | Q Exactive HF (Thermo Fisher Scientific) | LTQ Orbitrap Velos (Thermo Fisher Scientific) | LTQ Orbitrap (Thermo Fisher Scientific) |
| Data acquisition type | data-dependent acquisition | data-dependent acquisition | data-dependent acquisition |
| Software | KNIME workflow with PIA | MaxQuant 1.6.17.0 | MaxQuant 1.6.17.0 |
| Search engines | Mascot 2.7<br>MS-GF+<br>X!Tandem | Andromeda | Andromeda |
| Enzyme | Trypsin | Trypsin | Trypsin, LysC |
| Max. missed cleavages | 2 | 2 | 2 |
| Decoy generation | Randomize (concatenated target-decoy-db) | Revert | Randomize |
| Precursor tol. | 5 ppm | 20 ppm (first search)<br>6 ppm (main search) | 20 ppm (first search)<br>4.5 ppm (main search) |
| Fragment tol. | 20 mmu | 0.5 Da | 0.5 Da |
| Fixed PTMs | Carbamidomethyl (C) | Carbamidomethyl (C) | Carbamidomethyl (C) |
| Variable PTMs | Oxidation (M)<br>Gln-> pyro-Glu (N-terminal Q)<br>Deamidated (NQ)<br>Ammonium (DE)<br>Ammonialoss (N, N-terminal C) | Oxidation (M)<br>Acetyl (Protein N-term) | Oxidation (M)<br>Acetyl (Protein N-term) |

**S2 Table: O**verview over the quantitative peptide-level data sets.

| | D1_quant | D2_quant | D3_quant | D3_iso_quant |
|---|---|---|---|---|
| experimental groups | 5 | 9 | 2 | 2 |
| pairwise comparisons | 10 | 36 | 1 | 1 |
| quantified peptides* | 22,939 | 8,101 | 46,544 | 44,843 |
| mean number of peptides with valid ratio** | 18,093 | 5,619 | 30,369 | 29,490 |

* In at least one sample. Only peptides inside the chosen length range are considered.

** Mean over the pairwise group comparisons.

**S3 Table: I**nfluence of different minimal peptide lengths on the bipartite graphs for D2_fasta.

| | min 5 AA | min 6 AA | min 7 AA | min 9 AA |
|---|---|---|---|---|
| protein accessions | 6,336 | 6,335 | 6,333 | 6,333 |
| protein nodes | 6,265 | 6,264 | 6,263 | 6,263 |
| peptide sequences | 752,532 | 718,476 | 679,995 | 603,418 |
| peptide nodes | 12,962 | 8,132 | 7,563 | 7,366 |
| edges | 26,792 | 15,029 | 13,653 | 12,677 |
| graphs | 1,649 | 4,908 | 5,471 | 5,604 |
| graphs with 1 protein node | 1,506 | 4,296 | 5,045 | 5,264 |
| isomorphism classes | 18 | 64 | 41 | 36 |
| **largest graph*** | | | | |
| protein nodes | 4,413 | 116 | 69 | 69 |
| peptide nodes | 10,907 | 356 | 264 | 253 |
| edges | 24,434 | 1,292 | 3,329 | 3,033 |
| **second largest graph*** | | | | |
| protein nodes | 21 | 70 | 52 | 50 |
| peptide nodes | 37 | 267 | 203 | 181 |
| edges | 131 | 3,359 | 743 | 613 |

* In terms of number of protein nodes.

**S4 Table: I**nfluence of different minimal peptide lengths on the bipartite graphs for D3_fasta (without isoforms).

| | min 5 AA | min 6 AA | min 7 AA | min 9 AA |
|---|---|---|---|---|
| protein accessions | 81,591 | 81,572 | 81,548 | 81,440 |
| protein nodes | 80,932 | 80,897 | 80,856 | 80,676 |
| peptide sequences | 3,309,331 | 3,204,104 | 3,050,340 | 2,733,226 |
| peptide nodes | 192,162 | 157,556 | 148,555 | 143,264 |
| edges | 699,493 | 480,830 | 431,391 | 401,884 |
| graphs | 4,576 | 14,177 | 20,270 | 22,327 |
| graphs with 1 protein node | 3,722 | 8,178 | 10,129 | 11,088 |
| isomorphism classes | 253 | 2,305 | 4,198 | 4,522 |
| **largest graph*** | | | | |
| protein nodes | 74,157 | 40,266 | 6,472 | 2,203 |
| peptide nodes | 183,383 | 89,697 | 14,993 | 5,106 |
| edges | 685,438 | 315,173 | 56,950 | 17,897 |
| **second largest graph*** | | | | |
| protein nodes | 27 | 86 | 306 | 229 |
| peptide nodes | 51 | 126 | 757 | 454 |
| edges | 198 | 1,412 | 2,884 | 2,084 |

* In terms of number of protein nodes.

**S5 Table: C**omparison of bipartite graph characteristics without and with isoforms on data set D3 (with minimal peptide length of seven amino acids).

| | D3_fasta | D3_iso_fasta | D3_quant | D3_iso_quant |
|---|---|---|---|---|
| protein accessions | 81,548 | 103,541 | 17,585 | 22,216 |
| protein nodes | 80,856 | 102,463 | 10,969 | 11,672 |
| peptide sequences | 3,050,340 | 3,154,386 | 30,369 | 29,490 |
| peptide nodes | 148,555 | 184,932 | 10,540 | 10,895 |
| edges | 431,391 | 647,037 | 23,802 | 27,038 |
| graphs | 20,270 | 20,048 | 5,267 | 5,162 |
| graphs with 1 protein node | 10,129 | 8,948 | 3,315 | 3,106 |
| isomorphism classes | 4,198 | 5,657 | 459 | 524 |
| **largest graph*** | | | | |
| protein nodes | 6,472 | 8,895 | 57 | 58 |
| peptide nodes | 14,993 | 19,123 | 65 | 67 |
| edges | 56,950 | 87,852 | 350 | 359 |
| **second largest graph*** | | | | |
| protein nodes | 306 | 434 | 35 | 37 |
| peptide nodes | 757 | 940 | 34 | 34 |
| edges | 2,884 | 4,613 | 181 | 185 |

* In terms of number of protein nodes.

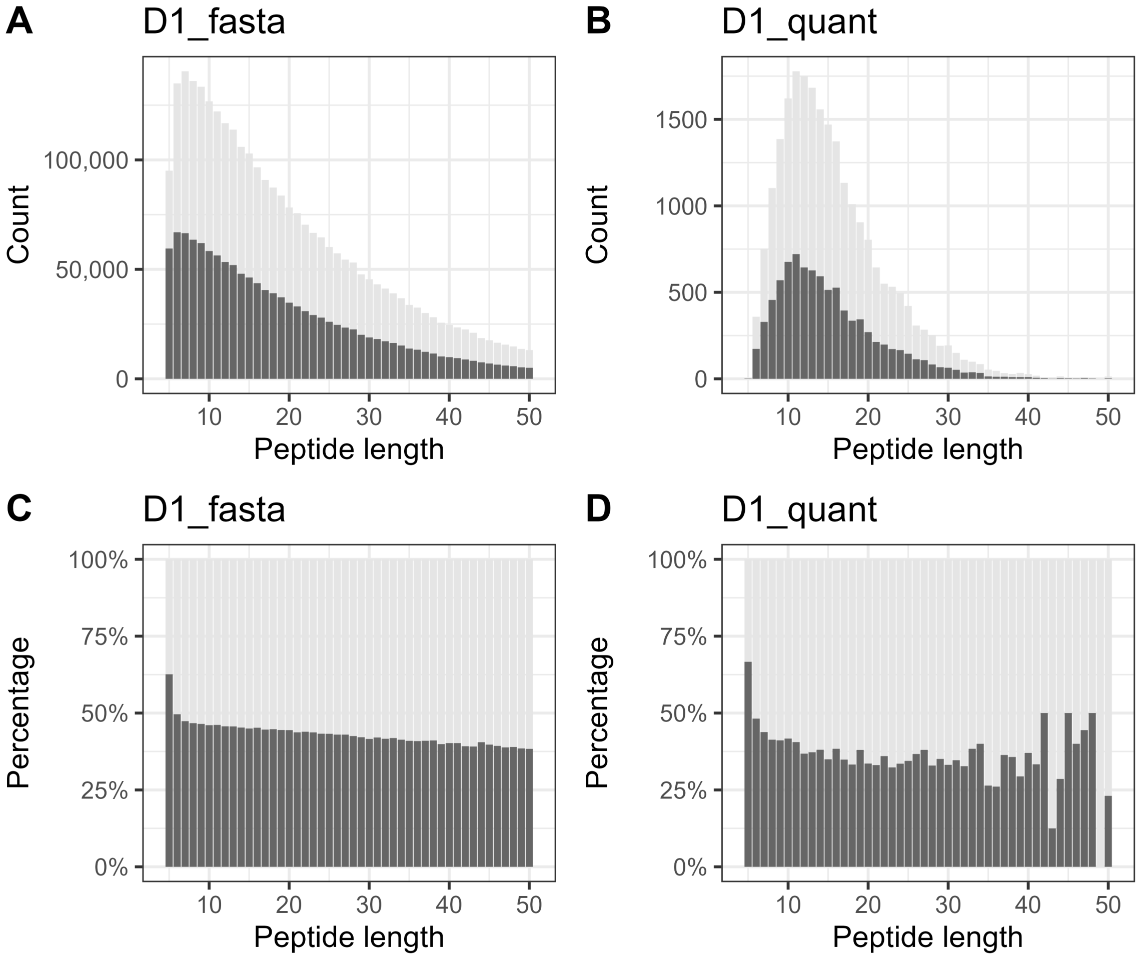


**S1 Figure: C**ount and percentages of shared and unique peptide sequences depending on the peptide length. The peptide length is given in amino acids. As an example, here the values for data set D1 are shown. Uniqueness is here defined as belonging to only one protein node, which may consist of multiple protein accessions.

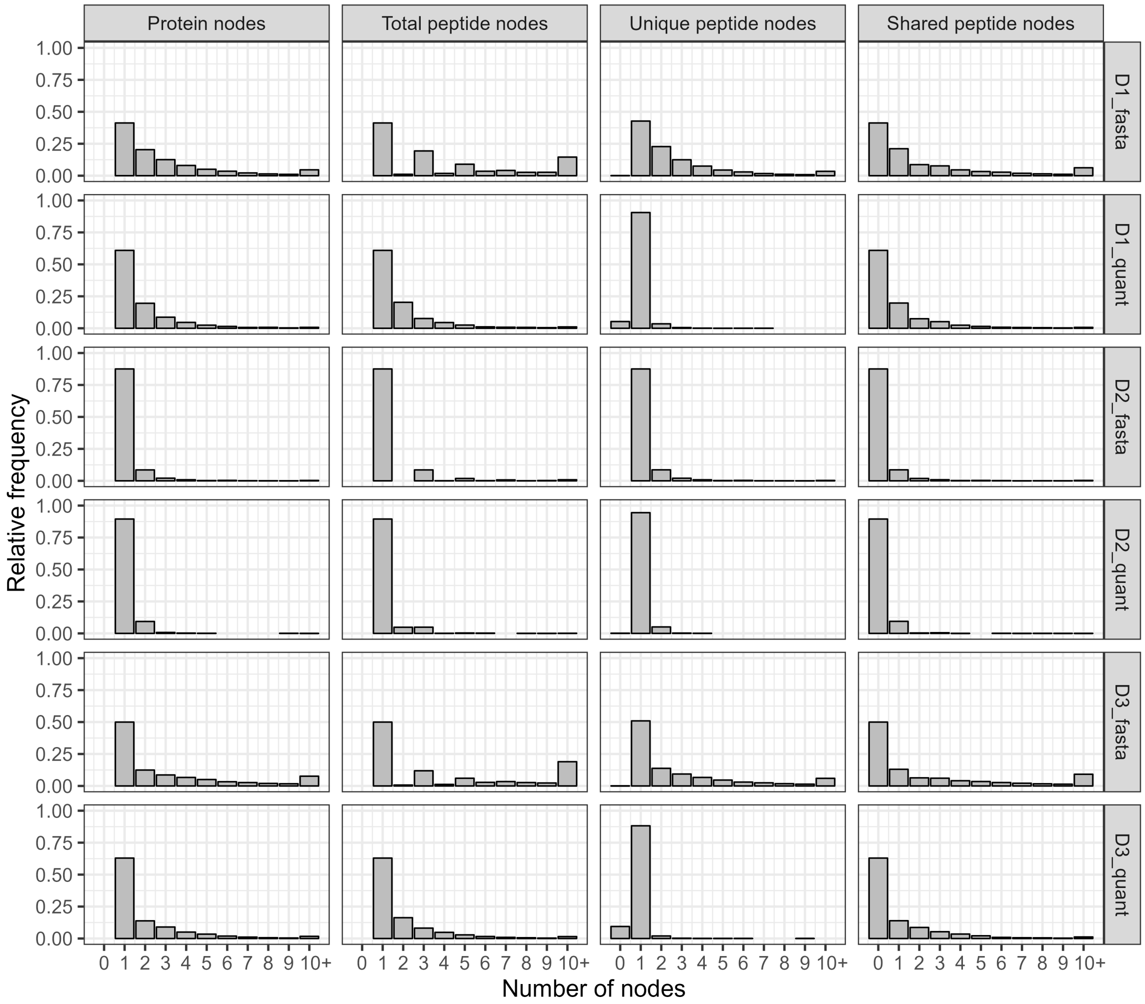


**S2 Figure: D**istribution of numbers of different node types for the bipartite peptide-protein graphs. The leftmost column shows the distribution of number of protein nodes for D1, D2 and D3, on database level (*__fasta) and on the level of quantified peptides (*__quant). On the x-axis the number of protein nodes are shown and on the y-axis the proportion of graphs with this exact number of protein nodes in comparison to all graphs. E.g., the bar at x = 1 is the percentage of graphs with only one protein node compared to all graphs. The sum of all bar heights adds up to one for each subfigure. Similarly, the distribution of the number of total peptide nodes, unique peptide nodes and shared peptide nodes are shown in the following columns. Minimal peptide lengths were chosen as seven for D1 and D3 and six for D2. The rightmost bar with the label ”10+” comprises the values of 10 and above.

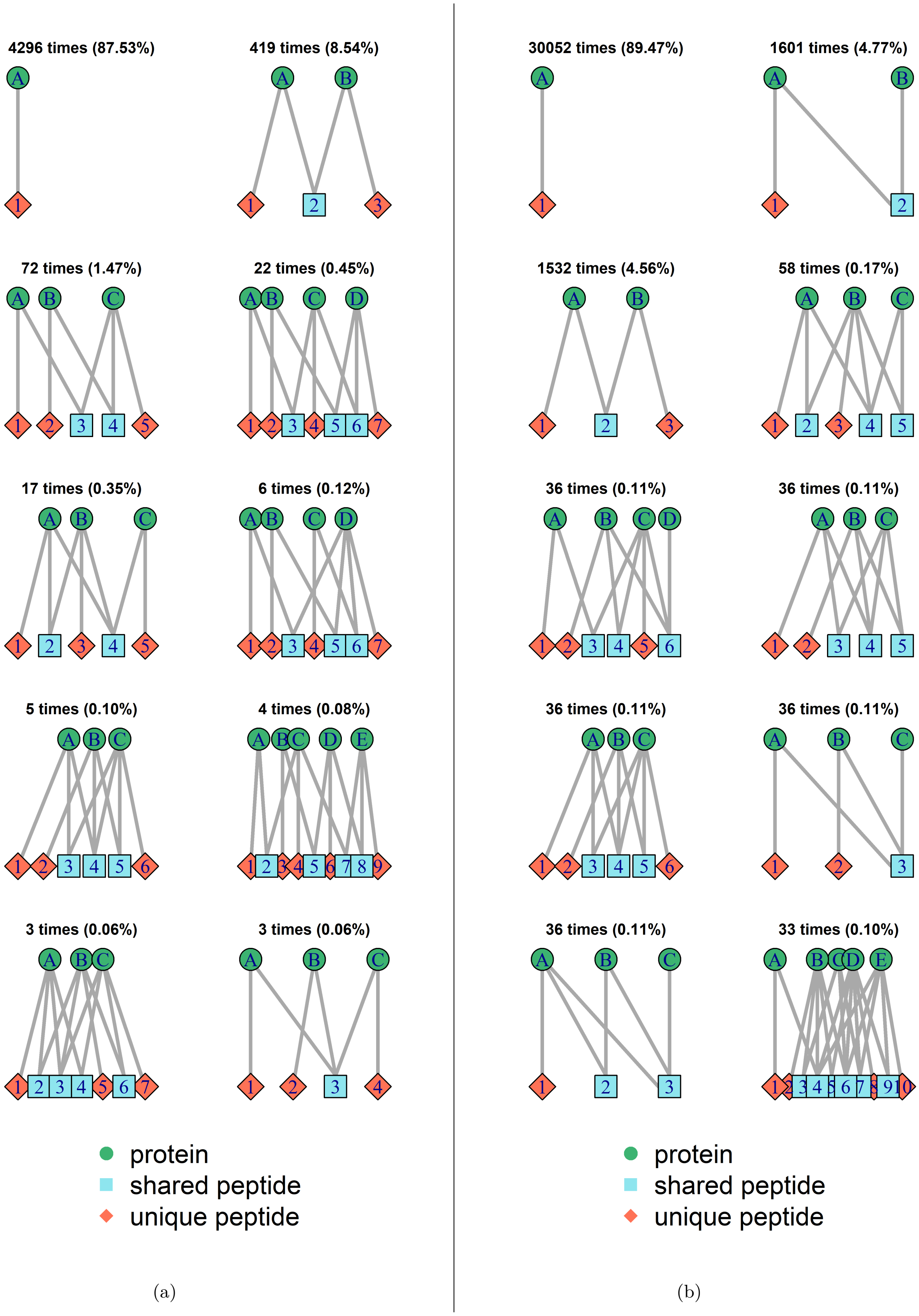


**S3 Figure: R**epresentative bipartite graphs of the ten largest isomorphism classes found in data set D2. (a) D2_fasta, (b) D2_quant, with number of occurrences and percentage of all graphs.

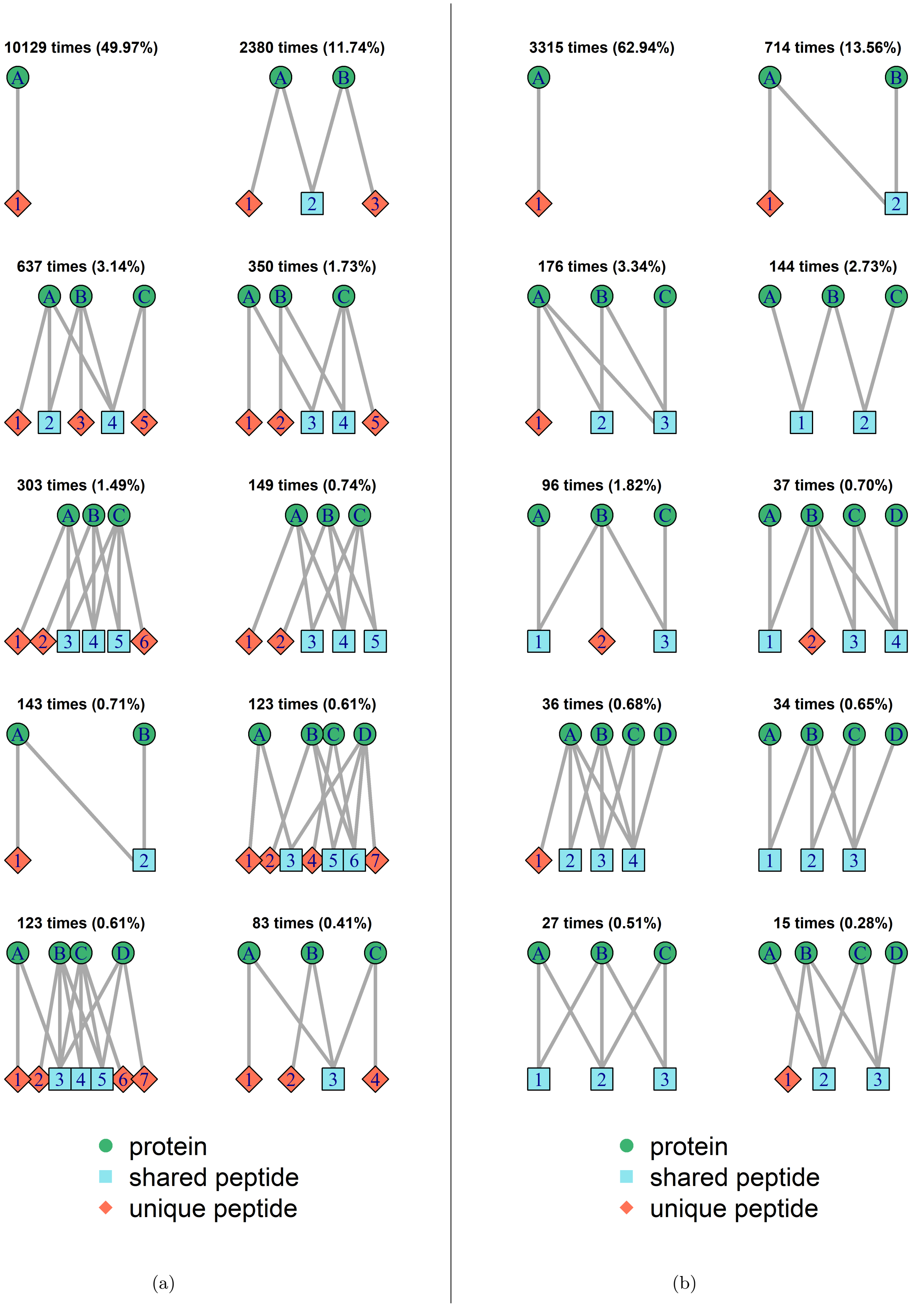


**S4 Figure: R**epresentative bipartite graphs of the ten largest isomorphism classes found in data set D3 (a) D3_fasta, (b) D3_quant, with number of occurrences and percentage of all graphs.

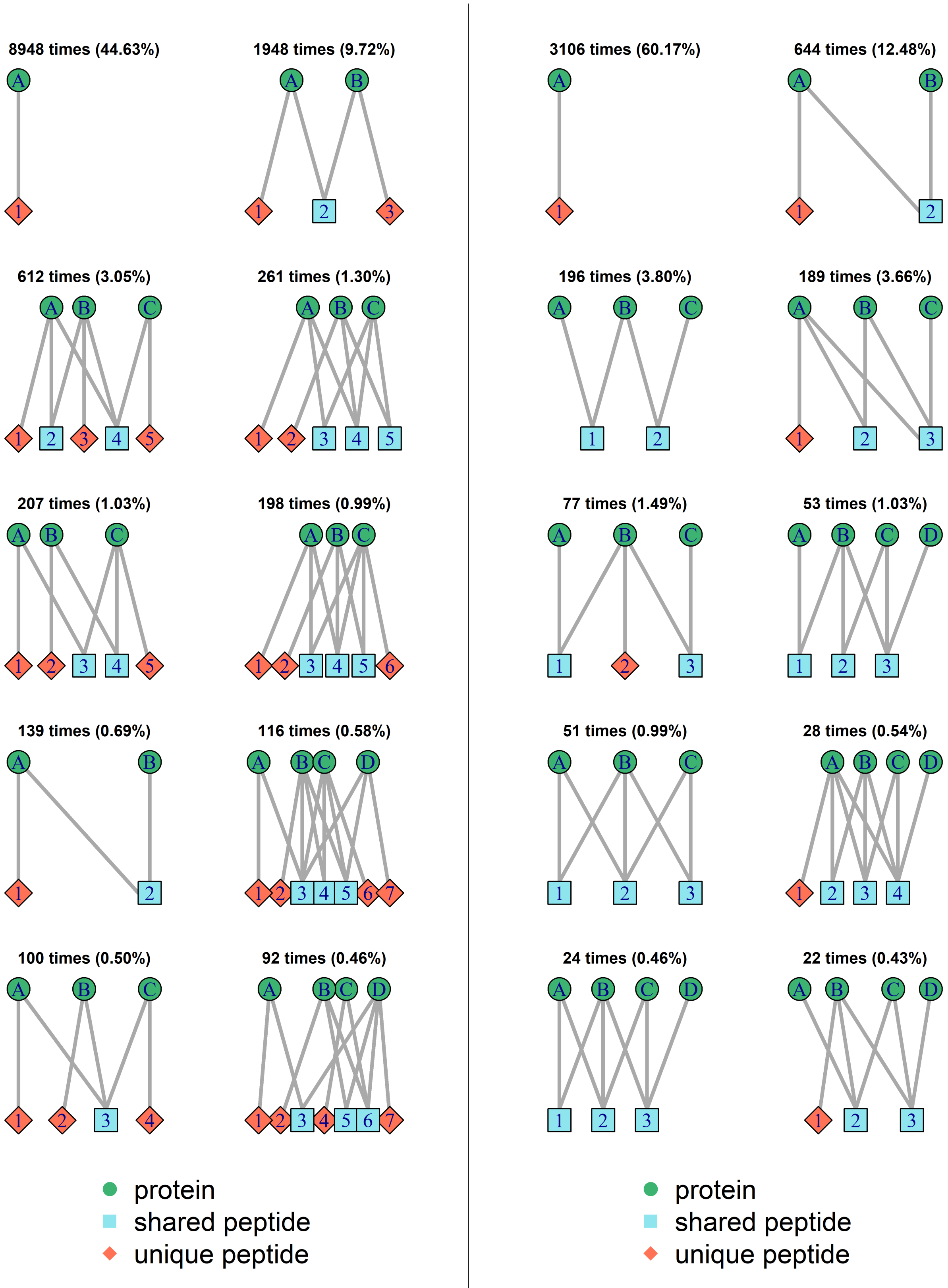


(a) (b)

**S5 Figure: R**epresentative bipartite graphs of the ten largest isomorphism classes found in data set D3_iso. (a) D3_iso_fasta, (b) D3_iso_quant, with number of occurrences and percentage of all graphs.
